# N-terminal modification of actin by acetylation and arginylation determines the architecture and assembly rate of linear and branched actin networks

**DOI:** 10.1101/2020.06.25.172320

**Authors:** Samantha M. Chin, Tomoyuki Hatano, Lavanya Sivashanmugam, Andrejus Suchenko, Anna S. Kashina, Mohan K. Balasubramanian, Silvia Jansen

## Abstract

The great diversity in actin network architectures and dynamics is exploited by cells to drive fundamental biological processes, including cell migration, endocytosis and cell division. While it is known that this versatility is the result of the many actin-remodeling activities of actin-binding proteins, recent work implicates post-translational modification of the actin N-terminus by either acetylation or arginylation itself as an equally important regulatory mechanism. However, the molecular mechanisms by which acetylation and arginylation alter the properties of actin are not well understood. Here, we directly compare how processing, and modification of the N-terminus of actin affects its intrinsic polymerization dynamics and its remodeling by actin-binding proteins that are essential for cell migration. We find that in comparison to acetylated actin, arginylated actin reduces intrinsic as well as formin-mediated elongation and Arp2/3-mediated nucleation. By contrast, there are no significant differences in Cofilin-mediated severing. Taken together, these results suggest that cells can employ the differently modified actins to precisely regulate actin dynamics. In addition, unprocessed, or non-acetylated actin show very different effects on formin-mediated-elongation, Arp2/3-mediated nucleation, and severing by Cofilin. Altogether, this study shows that the nature of the N-terminus of actin can induce distinct actin network dynamics, which can be differentially used by cells to locally finetune actin dynamics at distinct cellular locations, such as at the leading edge.

## Introduction

Actin cytoskeleton dynamics are the driving force for a myriad of essential cellular functions including cell migration, cytokinesis, intracellular transport, and contractility.^1, 2, 3^ The functional diversity of actin stems in part from its intrinsic ability to rapidly and reversibly transition between monomeric and filamentous forms, and is further expanded through tight spatiotemporal control by hundreds of actin-binding proteins with specific actin-remodeling activities.^4^ In addition, it is becoming increasingly clear that direct modification of actin molecules, for example by phosphorylation, acetylation, arginylation, methylation or oxidation, can have profound effects on actin network dynamics.^5, 6, 7, 8, 9, 10^

Posttranslational modification of the N-terminus of the abundant cytoplasmic β-actin by mutually exclusive acetylation or arginylation is emerging as a first-line mechanism to regulate cell migration.^11, 12, 13^ Like its cytoplasmic partner γ-actin, the N-terminal initiator Met of β-actin is removed cotranslationally, thereby exposing the second residue (Asp2 in β-actin, Glut2 in γ-actin) for further modification by acetylation (Arnesen et al., and references therein^14^). Although this extensive processing of the N-terminus of actin was already understood in the early ‘80s, it was not until recently that it was shown that acetylation of the exposed acidic residue is mediated by a dedicated N-acetyltransferase, NatH/NAA80, which specifically diverged to only target the acidic N-terminus of all six human actins.^15, 16^ Surprisingly, NAA80 KO cells are hypermotile and show significant increases in the number of filopodia and total F-actin content, whereas *in vitro* assays show that non-acetylated β/γ-actin displays slower filament elongation than acetylated β/γ-actin, even in the presence of formins.^12^ This leaves open the question of how acetylation of actin mechanistically contributes to curbing cell migration.

The N-terminus of the cytoplasmic actins can be further trimmed down to the second acidic residue, which then can be arginylated by the non-specific arginyltransferase Ate1.^7, 17^ Interestingly, arginylation in combination with slower translation leads to immediate proteasomal degradation of γ-actin.^18^ As a result, only arginylated β-actin (hereafter referred to as R-actin) is detected in cells, where it has been shown to specifically relocate to the leading edge upon induction of cell migration.^13^ Although less than 1% of the total β-actin population is estimated to be arginylated, it is possible that local concentrations at the leading edge during active cell migration are much higher and thus exert significant effects on local actin cytoskeleton dynamics.^11^ However, both actin as well as various actin-binding proteins can be modified by Ate1, making it difficult to identify what role and which actin-remodeling activities are directly influenced by arginylation of β-actin.^17^ Also, NAA80 KO cells show an 8-fold increase in R-actin, which supports the hypothesis that acetylation and arginylation of β-actin are mutually exclusive, and that the enhanced motility of NAA80 KO cells might at least in part be due to the increased presence of the more positively charged R-actin.^17^ This strongly suggests that acetylated (Ac-actin) and R-actin have different intrinsic dynamics and/or interact differently with key actin-regulatory proteins that control cell migration; however the underlying molecular mechanisms remain unclear to date.

Here, we have used pick-ya-actin, a recently established method to produce pure populations of specifically modified β-actin in Pichia Pastoris, to perform the first direct comparison between acetylated and arginylated mammalian β-actin.^19, 20^ To get a better understanding of the contribution of the charge at the N-terminus of β-actin, we also included pure populations of unprocessed (M-actin), and non-acetylated β-actin (D-actin). TIRF microscopy analysis elucidated that these actins have distinct intrinsic polymerization properties, and interact differently with key actin-binding proteins, including Profilin, mDia1, Arp2/3 and Cofilin. Altogether, this study shows for the first time that proper processing and different N-terminal modifications of actin can alter actin network dynamics and may thusly provide an extra layer of cytoskeletal regulation in cells.

## Results and Discussion

### Acetylation of actin markedly facilitates actin filament nucleation

The N-terminus of β-actin is cotranslationally processed to prepare it for further modification by either acetylation on Asp2 or arginylation on Asp3; however, it remains unclear how this extensive processing and subsequent modification, affect the intrinsic self-assembly dynamics of β-actin. To investigate this, we monitored polymerization of recombinantly produced Ac-actin, R-actin, unprocessed β-actin (M-actin), and non-acetylated β-actin (D-actin) in real time using TIRF microscopy (Fig. 1A). This method enables us to directly visualize and accurately quantify nucleation events, as well as measure filament elongation rates. Analysis of the intrinsic assembly behavior showed a striking difference in actin filament nucleation for Ac-actin, which was about 5-fold higher than any of the other actins (Fig. 1B-C and S1A). By contrast, individual actin filaments assembled from each actin were observed to elongate at similar rates (Fig. 1 D). Both Ac- and M-actin polymerized slightly faster than R- and D-actin, which is in agreement with an earlier study that compared acetylated versus non-acetylated actin.^12^ Contrary to our results, the same study did not report any difference in actin filament nucleation between Ac- and D-actin. However, this may be due to the fact that the actins used in prior work were isolated from WT or NAA80 KO HAP1 cells, and as a result consisted of a mixture of actin isoforms instead of pure β-actin. In addition, the actin in these studies was natively purified from mammalian cells, which makes it highly likely that their actin monomers might carry a multitude of other modifications that can have a compounding effect on their intrinsic nucleation behavior. However, despite the different experimental approaches, both studies show that complete processing and acetylation of the N-terminus can influence actin filament assembly. In addition, it demonstrates that there are clear differences in the intrinsic polymerization of R-actin and Ac-actin. The latter is in agreement with a study in *D. Discoideum,* which showed that actin isolated from Ate1 knockout cells polymerized slower than actin isolated from WT cells in bulk pyrene fluorescence assays.^21^

**Figure 1.**
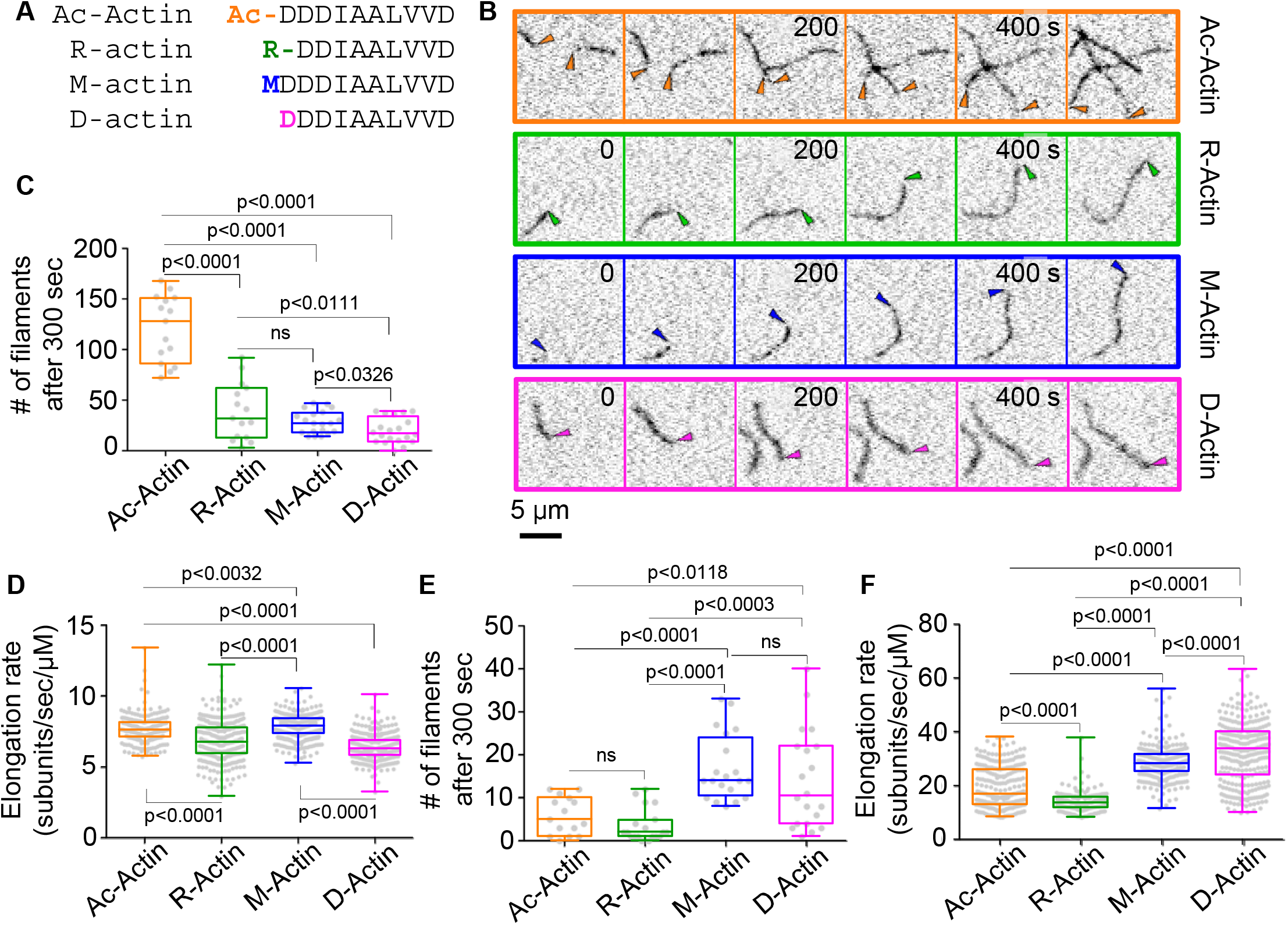
**(A)** Overview of the different actins used in this study. **(B)** Montages of representative TIRF microscopy movies showing the intrinsic self-assembly of the indicated actins. **(C, D)** Distribution of actin filament nucleation **(C)**, and filament elongation rate **(D)** of the differently modified actins. Corresponding box and whisker plots, indicating minimum, maximum and median, are shown as overlays. **(E, F)** Distribution of mDia1-mediated actin filament nucleation **(E)**, and elongation **(F)** of actins with different N-termini. Distributions shown in **C-F** are derived from at least three independent experiments with 5 FOVs per experiment. For nucleation, N ≥ 15 for each different actin. For elongation, N ≥ 182 individual filaments for each different actin. Corresponding box and whisker plots, indicating minimum, maximum and median, are shown as overlays. All P-values were obtained by comparing the indicated pairs of actins using an unpaired t-test.

### mDial can induce polymerization of both Ac- and R-actin, albeit at different rates

To further probe how modification of the N-terminus of actin contributes to actin filament assembly, we next investigated actin polymerization in the presence of profilin 1 (PFN) and the elongation-promoting formin, mDia1. These assays surprisingly showed a very different nucleation pattern for the differently modified actins compared to nucleation for each of these actins by themselves. First, Ac-actin was no longer the most efficient at nucleating actin filaments (Fig. 1E). Instead, nucleation by Ac-actin was similar to nucleation by R-actin, and about two-fold decreased compared to M-actin, and D-actin (Fig. 1E and S1B). These results show that PFN severely dampens nucleation of Ac-actin *in vitro* and may serve as a mechanism to curb the excessive and uncontrolled nucleation of Ac-actin in cells. In line with this hypothesis, it was recently shown that PFN increases the activity of the actin-specific acetyltransferase NAA80, suggesting that, in cells, Ac-actin is already sequestered by PFN even before it is modified.^16^ Together with our observation that D-actin nucleates better than Ac-actin in the presence of PFN (Fig. 1E), the latter also raises the question whether acetylation increases the affinity of PFN for Ac-actin.

In addition to nucleation, analysis of individual filaments elongated by mDia1 showed slower polymerization for both Ac-actin and R-actin as compared to M-actin and D-actin filaments (Fig. 1F and S1B). Further, polymerization by mDia1 using R-actin was slower compared to mDia1 elongation in the presence of Ac-actin, albeit only slightly (14.0 ± 0.3 subunits/sec/μM versus 19.3 ± 0.5 subunits/sec/μM). This demonstrates that formin-mediated elongation does not distinguish between Ac-actin and R-actin, and formins can use both actins as building blocks for filaments at the leading edge. However, it remains to be investigated whether the slightly slower elongation of R-actin by mDia1 also has a physiological significance. Our data further show that mDia1 polymerization is more effective in the presence of D-actin, which is the opposite of what was reported by Drazic and colleagues^12^. However, as pointed out above, their non-acetylated actin consisted of a mixture of actin isoforms containing native nonidentified modifications. In addition, it was demonstrated later that NAA80 KO cells contain up to 8-fold more R-actin.^11^ As such, non-acetylated actin purified from NAA80 KO cells also contains more R-actin, which, as we show here, displays a 2-fold lower polymerization rate than D-actin (Fig. 1F).

### Branched actin nucleation is reduced by arginylation of actin

R-actin shows a distinct lamellipodial localization in migrating cells^13^, suggesting that it participates in the formation of the extensive branched networks underneath the protruding cell membrane. In line with this, the branched actin network at the leading edge of Ate1 KO cells is severely disorganized, although it remains unclear whether this stems from arginylation of β-actin or the Arp2/3 complex, as both were identified as Ate1 targets.^17^ In addition, NAA80 KO cells show a marked increase in F-actin and the number of filopodia.^12^ This suggests that actin-binding proteins at the leading edge use acetylated and arginylated actin for different purposes, however, it is not clear how each modification alters the function of these proteins in regulating actin dynamics. Our approach with isoform-pure and singly modified actin uniquely enables us to investigate the direct effects of arginylation and acetylation of β-actin on branched actin network dynamics.

We first analyzed how each modification individually influences Arp2/3-mediated nucleation. For this purpose, we compared the formation of branches from the sides of filaments that were polymerized from either Ac-actin or R-actin. We observed that neither modification of the N-terminus abolished Arp2/3-mediated branch formation. In addition, we measured substantial differences in the total number of actin branches per field of view, demonstrating that Ac-actin produces almost 7-fold more branches than R-actin (Fig. S2A). However, to nucleate a new daughter filament the Arp2/3 complex needs a mother filament. Consequently, the number of daughter filaments should increase with the number of available mother filaments, and thus actin filament nucleation, which we showed to be greatly enhanced for Ac-actin (Fig. 1C). To ensure that the densely arborized networks obtained with Ac-actin are due to an effect on Arp2/3 nucleation, we accounted for its enhanced intrinsic nucleation by determining the number of branches that occurred per micron actin filaments, as well as by following the increase in the number of branches over time. These analyses clearly show that both branch density (Fig. 2B) and branching rate (Fig. 2C and S2B) are significantly higher for Ac-actin than R-actin. In addition, this experiment also demonstrated that M-actin greatly increased Arp2/3 branching density, without a concomitant increase in branching rate, whereas D-actin showed a reduced branch density and branching rate compared to Ac-actin (Fig. 2B-C). Altogether, this suggests that N-terminal acetylation or arginylation of actin can regulate the spacing and number of actin branches, although it remains elusive whether this is due to a direct effect on the Arp2/3 complex or the activating nucleation-promoting factors (NPFs).

**Figure 2.**
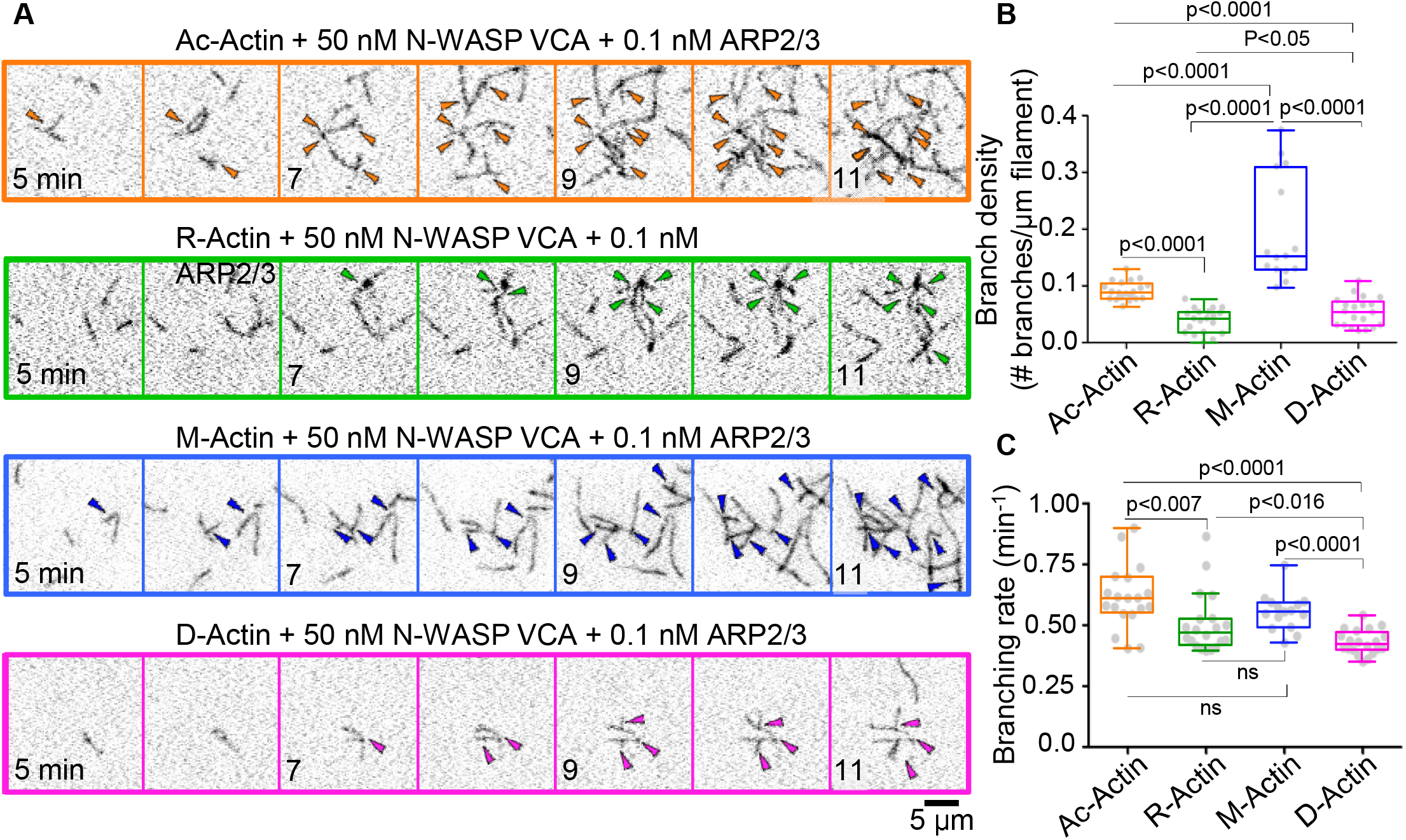
**(A)** Montages of representative TIRF microscopy movies. Arrowheads indicate newly formed actin branches. **(B, C)** Distribution of branch density **(B)** and branching rates **(C)** of the different actins (N ≥ 15, 3 or 4 replicates with 5 FOV each). Corresponding box and whisker plots, indicating minimum, maximum and median, are shown as overlays. P-values were obtained by comparing the indicated actins using an unpaired t-test.

Our analysis shows that branched actin networks are assembled differently from pure populations of Ac-actin, and R-actin. However, in cells, both Ac-actin and R-actin localize to the leading edge, which raises the question how branched actin assembly is regulated in the presence of both modified actins. To investigate this, we prepared labeled R-actin, and followed its polymerization (10% labeled final) in the absence, as well as the presence, of increasing amounts of Ac-actin. These assays showed that mixing Ac-actin with R-actin increases total actin polymer mass (Fig. 3A-B), which is in agreement with the increased nucleation and elongation we observed for Ac-actin in this study (Fig. 1C-D). Surprisingly, we did not detect a similar effect for actin filament branching. Here, our analysis showed that addition of Ac-actin to R-actin did not noticeably alter the branching rate (Fig. 3C), even when the reaction consisted mostly of Ac-actin (30% R-actin, 70% Ac-actin). This suggests that in the presence of both actins, Arp2/3-mediated nucleation is still more sensitive to R-actin. We hypothesize that is either due to R-actin monomers that are incorporated into the mother filament or to R-actin monomers being bound preferentially by the NPFs. The former would affect the binding of Arp2/3 to the sides of existing filaments, whereas the latter would alter activation of Arp2/3-mediated nucleation of daughter filaments by NPFs.

**Figure 3.**
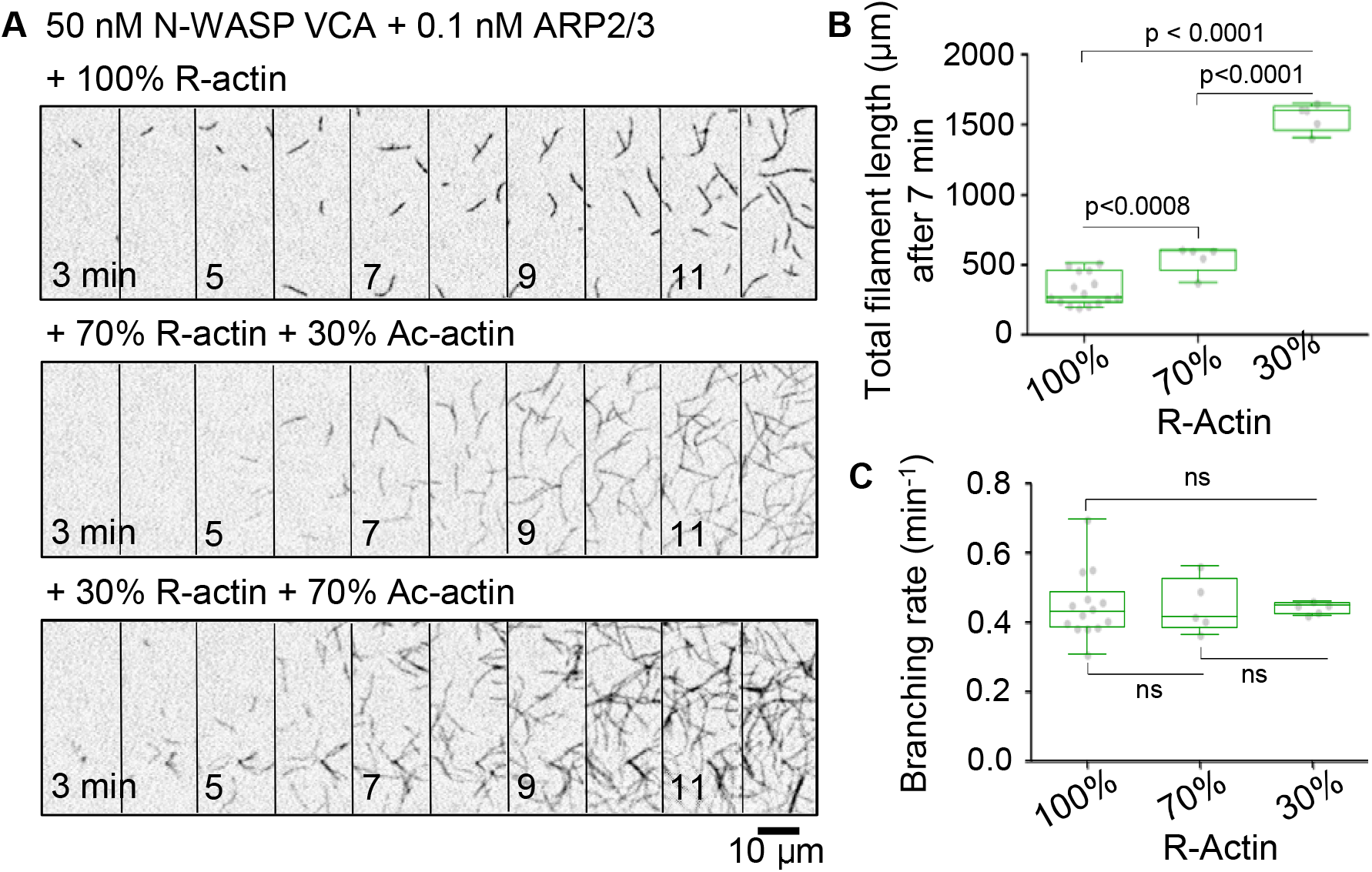
**(A)** Montages of representative TIRF microscopy movies. **(B, C)** Distribution of total filament length **(B)** and branching rates **(C)** of reactions consisting of different R-actin and Ac-actin ratios (N ≥ 5, 1 to 3 replicates with 5 FOV each). Corresponding box and whisker plots, indicating minimum, maximum and median, are shown as overlays. P-values were obtained by comparing the indicated ratios using an unpaired t-test.

### N-terminal modification has no effect on Cofilin-mediated severing

Branched actin network turnover depends on the formation of new branches by the Arp2/3 complex, as well as on continuous disassembly of older parts of the network, which is largely regulated by the filament severing protein, Cofilin. To assess whether acetylation and arginylation also affect branched network disassembly, we directly compared Cofilin-mediated severing of Ac-actin and R-actin filaments using TIRF microscopy. Our results show a small reduction in the cumulative severing rate of R-actin compared to Ac-actin, however this slight decrease did not significantly change the time to half-maximal severing (Fig. 4B-C, S3). Together with our observations that Ac-actin and R-actin differentially impact Arp2/3-mediated nucleation, and formin-mediated elongation, this strongly suggests that these differently modified actins are more likely to play a role in the assembly of different actin structures than in regulating actin network turnover.

**Figure 4.**
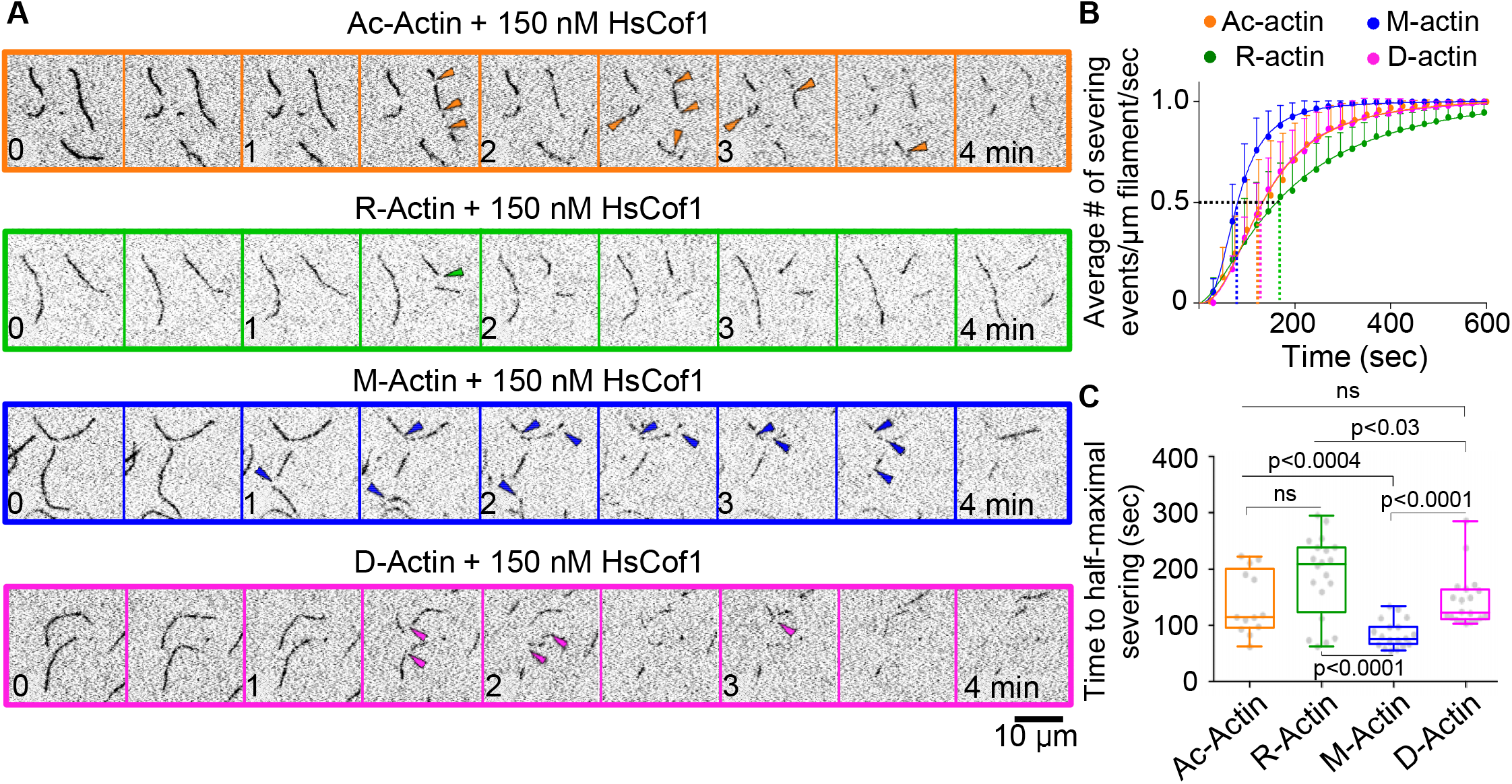
**(A)** Montages of representative TIRF microscopy movies. Arrowheads indicate severing events. **(B)** Cumulative severing of the indicated modified actin shown as the average ± StDev of the cumulative severing curves shown in Fig S3 (N = 9, 3 replicates with ≥ 3 FOV each). **(C)** Distribution of time to half-maximal severing calculated from the cumulative severing curves shown in Fig S3 (N = 9, 3 replicates with ≥ 3 FOV each). Corresponding box and whisker plots are shown as overlays. Statistical significance was tested by comparing two types of actin using an unpaired t-test.

Interestingly, we did not observe any difference in Cofilin-mediated severing between Ac-actin and D-actin. Taken together with the increased formin-mediated polymerization of D-actin (Fig. 1F), this suggests that the increase in filopodia observed in the absence of N-acetylation of actin are more likely due to altered actin assembly than disassembly kinetics.^12^ In addition, fragmentation of unprocessed M-actin was clearly enhanced in comparison to any of the other types of actins, suggesting that proper modification of the N-terminus results in a filament that is more resistant to the conformational changes induced by Cofilin, and thus is less prone to fragmentation. This again adds to our conclusion that N-terminal modification of actin rather affects actin network assembly than disassembly.

## Conclusions

It has been known for many years that actin monomers and filaments can undergo a multitude of modifications, however the functional impact of many of these modifications remains elusive. The fact that we can now produce populations of specifically modified actins enables us to start investigating how these modifications contribute to actin cytoskeleton dynamics. Here, we focus on directly comparing the effect of arginylation and acetylation of the N-terminus of β-actin on actin-binding proteins that shape the actin networks at the leading edge. Our results show that Ac-actin and R-actin differ most in intrinsic nucleation (Fig. 1B-C), however this effect is abolished in the presence of PFN (Fig. 1E). In comparison to Ac-actin, R-actin shows reduced elongation by mDia1 (Fig. 1F), and nucleation by Arp2/3 (Fig. 2). The latter persists even in the presence of substantial amounts of Ac-actin (Fig. 3). Based on our results, we postulate that Ac-actin and R-actin co-assemble into linear and branched actin networks at the leading edge, but that the ratio of R- to Ac-actin determines the rate at which these actin networks are being formed, as well as their final architecture (Fig. 5). In line with this, R-actin levels increase when cell migration is induced, suggesting that the specific R-actin branching rate is part of the cellular mechanism that drives robust protrusion.^13^ Our study further raises the question whether cells distinguish between these modified actins in their monomeric or filamentous forms. To address this, future studies will have to investigate in detail whether actin monomer binding proteins, such as PFN, Ena/VASP and NPFs, show a higher affinity for specifically modified actin monomers, and how this affects their cellular activities. In line with this, a recent study showed that actin filament elongation by the formin, INF2, is inhibited by a complex of Srv2/CAP and lysine-acetylated actin.^5, 22^ In addition, future work should analyze whether N-terminal modification of actin by acetylation and arginylation alters the intrinsic interactions of adjacent protomers in the actin filament, interaction of formins with the barbed end, or binding of the Arp2/3 complex to the side of mother filaments. Finally, we only tested a handful of actin-binding proteins, and more work is needed to gain insight into how these modifications influence other actin-remodeling activities, including F-actin crosslinking, bundling, stabilization, and disassembly.

**Figure 5.**
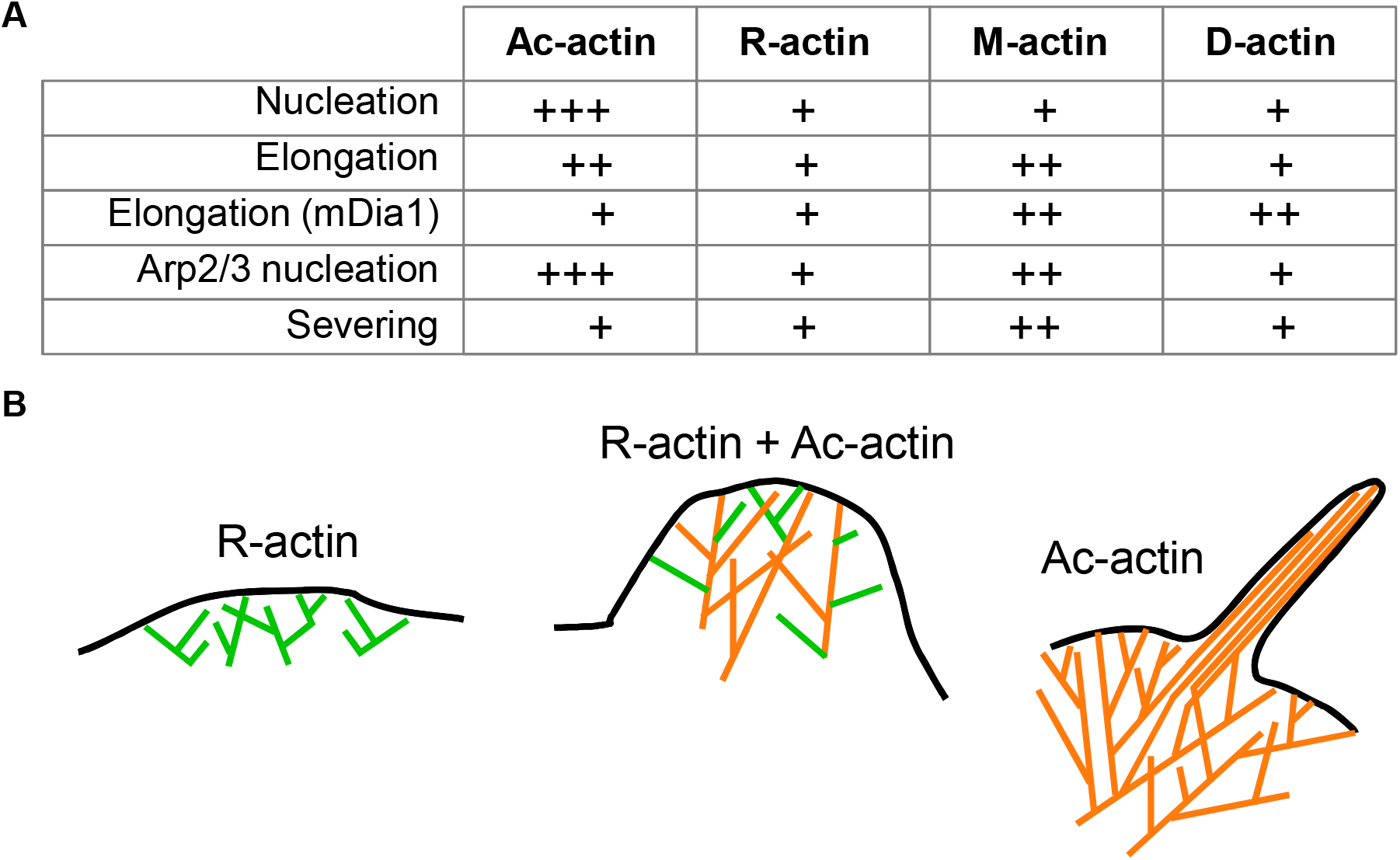
**(A)** Table comparing the effects of actins with different N-termini on the indicated actin-remodeling activities. **(B)** Model depicting the branched actin networks assembled by R-actin and Ac-actin. As both actins can be used for Arp2/3 nucleation, and formin-mediated elongation, we hypothesize that they co-assemble in branched and linear actin networks with different dynamics, the latter which is determined in part by the ratio of R- to Ac-actin, and the distinct effects these actins have on filament polymerization.

## Acknowledgments

The work in the laboratory of MBK was supported by a Wellcome Trust Collaborative Award in Science (203276/Z/16/Z), a European Research Council Advanced Grant (ERC-2014-ADG No. 671083), and a collaborative BBSRC award to MKB (BB/S003789/1). The work in the AK laboratory was supported by NIH grants R35GM122505 and R01NS102435.

## Materials and Methods

### Protein expression and purification

#### N-terminally modified Ac-β-actin, R-β-actin, M-β-actin, and D-β-actin

*P. pastoris* transformants for Ac-β-actin, R-β-actin, M-β-actin, and D-β-actin stored at −80°C were revived on YPD solid media plates at 30°C. Cells were inoculated into 200 ml MGY liquid media composed of 1.34% yeast nitrogen base without amino acids (SIGMA Y0626), 0.4 mg/L biotin and 1% glycerol and cultured at 30°C, 220 rpm. The culture medium was diluted to 6 L with fresh MGY media and cells were further cultured at 30°C, 220 rpm in six 2 L flasks until the optical density at 600 nm (OD_600_) reached around 1.5. Cells were pelleted down by centrifugation (10628 g at 25°C for 5 min, Thermo Fisher SCIENTIFIC #F9-6 x 1000 LEX rotor). The cells were washed once with sterilized water and re-suspended into 6 L MM composed of 1.34% yeast nitrogen base without amino acids (SIGMA Y0626), 0.4 mg/L biotin and 0.5% methanol. Cells were cultured in twelve 2 L baffled flasks (500 mL for each) at 30°C, 220 rpm for 1.5-2 days. 0.5 % methanol was fed every 24 hours during the culture. Cells were pelleted down by centrifugation (10628 g at 25°C for 5 min, Thermo Fisher SCIENTIFIC #F9-6 x 1000 LEX rotor). Cells were washed once with water and suspended into 75 mL ice cold water. The suspension was dripped into a liquid nitrogen bath and stored at −80°C. 50 g cell suspension was loaded into a grinder tube (#6801, SPEX^®^ SamplePrep) pre-cooled with liquid nitrogen. Cells were ground with freezer mill (#6870, SPEX^®^ SamplePrep) in a liquid nitrogen bath. Duration of the grinding was 1 min with 14 cycle per second (cps). The grinding procedure was repeated 30 times with 1 min intervals. Liquid nitrogen was re-filled every 10 times of the grinding. The resulting powder was kept on dry ice until the next step. The powder was thawed and resuspended in equal amounts of 2 x Binding buffer (20 mM imidazole (pH 7.4), 20 mM HEPES (pH 7.4), 600 mM NaCl, 4 mM MgCl_2_, 2 mM ATP (pH 7.0), 2x concentration of protease inhibitor cocktail (cOmplete, EDTA free #05056489001, Roche), 1 mM phenylmethylsulfonyl fluoride (PMSF) and 7 mM beta-mercaptoethanol). The lysate was sonicated on ice (3 minutes, 5 seconds pulse, 10 seconds pause with 60% amplitude, QSONICA SONICATORS) until all aggregates were resolved. The lysate was centrifuged at 4°C (3220 g for 5 min, Eppendorf #A-4-81 rotor) to remove intact cells and debris. Insoluble fraction was removed by high speed centrifugation at 4°C (25658 g for 30 minutes, Thermo Fisher SCIENTIFIC #A23-6 x 100 rotor). The supernatant was filtrated with 0.22 μm filter (Filtropur BT50 0.2, 500 ml Bottle Top Filter, #83.1823.101) and incubated with 6 ml Nickel resin (Thermo SCIENTIFIC #88222) at 4°C for 1 hour. The resin was pelleted down by centrifugation at 4°C (1258 g for 5 min, Eppendorf #A-4-81 rotor) and washed with ice-cold 50 mL Binding buffer composed of 10 mM imidazole (pH 7.4), 10 mM HEPES (pH 7.4), 300 mM NaCl, 2 mM MgCl_2_, 1 mM ATP (pH 7.0) and 7 mM betamercaptoethanol for 4 times. The resin was washed 5 times with G-buffer composed of 5 mM HEPES (pH 7.4), 0.2 mM CaCl_2_, 0.01 w/v% NaN_3_, 0.2 mM ATP (pH 7.0) and 0.5 mM dithiothreitol (DTT). The resin was suspended into ice-cold 40 mL G-buffer with 5 μg/ml TLCK treated chymotrypsin (SIGMA #C3142-25MG) and incubated overnight at 4°C. The chymotrypsin was inactivated by 1 mM PMSF and the elution was collected into a tube. Actin retained on the resin was eluted with 12 mL G-buffer (without DTT for labelling) and all elution fractions were combined. The eluent with R-β-actin was divided to perform labeling of half of the protein. Each portion was concentrated with 30 kDa cut-off membrane (SIGMA-ALDRICH #Z677892-24EA) to 0.9 ml and 2.5 ml. Ac-β-actin, M-β-actin, and D-β-actin were not labeled and hence the eluent of each actin was concentrated to 0.9 ml. The 0.9 ml of each actin was polymerized by adding 100 μl of 10x MKE solution composed of 20 mM MgCl_2_, 50 mM ethylene glycol tetraacetic acid (EGTA) and 1 M KCl for 1 hour at room temperature and was kept unlabeled. The 2.5 ml of R-β-actin sample was desalted to remove DTT leftovers by using PD-10 (Prepacked Disposable PD-10 Columns, GE Healthcare #17085101) column pre-equilibrated with G-buffer **without DTT**. Then 100 mM KCl and 2 mM MgCl_2_ were added into the desalted actin solution and the sample was rotated in cold room for 1 hour to induce actin polymerization. Next 3-fold excess of Alexa Fluor 647 C2 Maleimide (Thermo Fisher Scientific #A20347) was mixed with the solution and the mixture was incubated at room temperature for 1 hour. The cysteine-maleimide reaction was quenched by addition of 10 mM DTT at 4°C. The solution was centrifuged at ~3220 g for 5 min (Eppendorf #A-4-81 rotor) to pellet down precipitated dyes. The labeled and unlabeled polymerized actin samples were pelleted down by ultracentrifugation at room temperature (45000 rpm for 1 hour, Beckman TLA-55 rotor). The pellets were rinsed once with 1 mL G-buffer and re-suspended into ice cold 0.5 mL G-buffer. The actin was depolymerized by dialysis against 1 L G-buffer at 4°C for 2 days. The dialysis buffer was exchanged every 12 hours. The solutions with the depolymerized labeled and unlabeled actin were collected into 1.5 ml centrifuge tubes and stored on ice.

#### RMA and labeled RMA

RMA was purified by generating acetone powder from ground muscle tissue (Graziano et al., 2013), which was stored in aliquots at −80°C. To extract G-actin, one aliquot of acetone powder was pulverized using a coffee grinder, resuspended in G-buffer, and cleared by low-speed centrifugation and filtration through Whatman paper. The actin was allowed to polymerize overnight and then pelleted. The pellet was disrupted by dounce homogenization and dialyzed against G-buffer for 2 days with fresh G-buffer exchanges every day. To label actin with NHS-LC-LC-biotin (ThermoScientific) or AlexaFluor488-maleimide (ThermoFisher), G-actin was dialyzed against G-buffer (pH8.0) or G-buffer without DTT, respectively. Next, G-actin was polymerized in the presence of 3-fold excess of NHS-LC-LC-biotin or 5-fold excess of Alexa Fluor 488 maleimide for 2 hours at room temperature or overnight at 4°C. Labeled F-actin was collected by centrifugation. The labeled F-actin pellets were disrupted by dounce homogenization, dialyzed against G-buffer for 3 days with fresh G-buffer exchanges every day, and gel filtered on a 16/60 S200 column (GE Healthcare). Peak elution fractions of biotin-actin were combined and stored at −80°C after determination of the stock concentration. Combined peak fractions of Alexa488-actin were dialyzed to G-buffer with 50% glycerol, and stored at −20°C. Labeling efficiency of Alexa488-actin was measured by absorbance at 290 and 494 nm, and an extinction coefficient of 72,000 M^-1^cm^-1^. The absorption at 290 nm was corrected for background fluorescence from the AlexaFluor488 dye (correction factor 0.138).

#### Actin-regulatory proteins

pTYB11-mDia1(FH1-FH2), and pTYB11-HsPFN1 were kindly provided by Dr. R. Dominguez (UPenn, PA). pET15b-HsCof1, and pGAT2 GST-VCA (421-505) Hs N-WASP were obtained from Dr. Bruce Goode (Brandeis University, MA). All aforementioned proteins were expressed in Rosetta (DE3) *E. coli* by growing cells at 37°C in TB medium to log phase, then inducing expression with 0.5 mM isopropyl β-d-1-thiogalactopyranoside (IPTG) at 18°C for 16 h. Cells were harvested by centrifugation and stored at −80°C until purification.

For purification of pTYB11-mDia1(FH1-FH2), and pTYB11-HsPFN1, the frozen pellet was resuspended in 20 mM Tris (pH 7.4), 500 mM NaCl, 1 mM EDTA, protease inhibitor mix (Roche), 1mM PMSF, and 4 mM benzamidine. Next, the cells were lysed by three passes through a microfluidizer and cleared by centrifugation at 30,000 × *g* for 20 min in a Fiberlite F13-14 × 50CY rotor (ThermoScientific, Rockport, IL). The cleared lysate was passed over chitin resin (New England Biolabs, MA) by gravity, and the resin was washed with at least 10 column volumes of 20 mM Tris (pH 7.4), 500 mM NaCl, 1 mM EDTA. Bound proteins were eluted after 24h incubation in 20 mM Tris (pH 7.4), 500 mM NaCl, and 50 mM DTT to induce self-cleavage of the intein tag. Eluted mDia1(FH1-FH2), and HsPFN1 were further purified by gel filtration on a 16/60 S200 or 16/60 S75 column (GE Healthcare), respectively, equilibrated in 20 mM Tris (pH 7.4), 50 mM KCl, and 1 mM DTT. Peak fractions were concentrated, snap-frozen in liquid N_2_, and stored at −80°C until use. Protein concentration was determined spectrophotometrically by measuring the absorbance at 280 nm, and using calculated extinction coefficients of 21430 M^-1^cm^-1^ for mDia1(FH1-FH2), and 18450 M^-1^cm^-1^ for HsPFN.

For purification of pET15b-HsCof1, the frozen pellet was resuspended in 3-fold volume of 20 mM Tris (pH 8.0), 50 mM NaCl, 1 mM DTT, protease inhibitor mix, 1mM PMSF, and 4 mM benzamidine. Next, the bacterial cells were lysed by three passes through a microfluidizer, and cleared by centrifugation at 30,000 × *g* for 20 min. The cleared lysate was applied to a 5 ml HiTrap HP Q column (GE Healthcare Biosciences). The flow-through, containing HsCof1, was collected and dialyzed to 20 mM HEPES (pH 6.8), 25 mM NaCl, and 1 mM DTT. Next, the protein was applied to a 5 ml HiTrap SP FF column (GE Healthcare Biosciences) and eluted with a linear gradient of NaCl (25-500 mM). Fractions containing HsCof1 were concentrated and dialyzed to 20 mM Tris (pH 7.4), 50 mM KCl, and 1 mM DTT, aliquoted, snap-frozen in liquid N_2_, and stored at −80°C until use. Protein concentration was determined spectrophotometrically by measuring the absorbance at 280 nm, and using a calculated extinction coefficient of 14,440 M^-1^cm^-1^.

For purification of pGAT2 GST-VCA (421-505) HsN-WASP, the frozen pellet was resuspended in 3-fold volume of 1X PBS (137 mM NaCl, 2.7 mM KCl, 10 mM Na_2_HPO_4_, 1.8 mM KH_2_PO_4_), protease inhibitor mix, 1mM PMSF, and 4 mM benzamidine. Next, the bacterial cells were lysed by three passes through a microfluidizer, and cleared by centrifugation at 30,000 × *g* for 20 min. The cleared lysate was passed over glutathione sepharose 4B resin (GE heathcare) by gravity, and the resin was washed with at least 10 column volumes of 1X PBS. Bound proteins were eluted by 1X PBS with 10 mM reduced glutathione (Sigma Aldrich). Fractions containing VCA-NWASP were concentrated and dialyzed to 20 mM Tris (pH 7.4), 150 mM NaCl, and 1 mM DTT, aliquoted, snap-frozen in liquid N_2_, and stored at −80°C until use. Protein concentration was determined spectrophotometrically by measuring the absorbance at 280 nm, and using a calculated extinction coefficient of 5500 M^-1^cm^-1^.

Arp2/3 complex was purified from cryoground bovine brain (Pel-freez, Arkansas) using a modified protocol based on Boczkowska, et al (2008). Briefly, 200 g of frozen brain powder was resuspended in 400 ml of resuspension buffer (20 mM Tris (pH 8.0), 120 mM NaCl, 5 mM MgCl_2_, 5 mM EGTA, 1 mM DTT) with protease inhibitor mix, 1 mM PMSF, and 4 mM benzamidine, and clarified by centrifugation at 12,000 × g for 30 min. The cleared supernatant was loaded onto Q - sepharose Fastflow resin (GE Healthcare) pre-equilibrated with resuspension buffer. The flow-through, containing Arp2/3 complex, was applied onto a VCA-HsNWASP affinity column equilibrated with resuspension buffer supplemented with 0.1 mM ATP. Bound Arp2/3 complex was eluted in 20 mM Tris (pH 8.0), 25 mM KCl, 400 mM MgCl_2_, 1 mM EGTA, 1 mM DTT, 0.1 mM ATP, and dialyzed against 20 mM MES (pH 6.4), 20 mM KCl, 2 mM MgCl_2_, 1 mM EGTA, and 0.1 mM ATP, before further purification on a Mono S column (GE Healthcare) using a 20 mM −1M KCl gradient. Peak fractions were combined, and dialyzed against 20mM Tris HCl (pH 7.5), 50mM KCl, 2mM MgCl_2_, 1mM DTT, 10% glycerol, 1mM DTT, 0.1 mM ATP, aliquoted, snap-frozen in liquid N_2_, and stored at −80°C until use. The concentration of the purified complex was determined spectrophotometrically using a theoretically calculated extinction coefficient of 234,080 M^-1^ cm^-1^.

### TIRF microscopy (TIRFM)

Prior to experiments, all actins were dialyzed overnight against G-buffer (3 mM Tris pH 7.5, 0.5 mM DTT, 0.2 mM ATP pH 7.0, and 0.1 mM CaCl_2_). Next day, actins were cleared by highspeed centrifugation at 316,613 x g for 1 hour at 4°C in a TLA100 rotor (Beckman Coulter). Only the top 80% of the supernatant was collected to perform TIRF microscopy. Labeling efficiency and concentration was determined for each actin after centrifugation using a DS-11 + spectrophotometer (Denovix). Labeling of Alexa-488-actin was calculated by measuring absorbance at 290 and 494 nm using the extinction coefficient of 72000 M^-1^ cm^-1^, and a Alexa488 correction factor of 0.138. Labeling efficiency of ATTO-560 labeled actin was determined by measuring absorbance at 290 and 554 nm using an extinction coefficient of 120000 M^-1^ cm^-1^, and a correction factor of 0.12.

For all TIRFM experiments, coverslips were sonicated for 1 hour in detergent, 20 min in 1 M KOH, 20 min in 1 M HCl, and 1 hour in 100% ethanol. Clean coverslips were then rinsed with ddH_2_O, dried extensively in an N_2_-stream, and coated with 200 μl/slide of freshly prepared coating solution (80% ethanol pH 2.0, 2 mg/ml methoxy-(ethylene glycol)-silane, and 4 μg/ml biotin-poly(ethylene glycol-silane)). Coated coverslips were incubated overnight at 70°C for 16 hours, rinsed extensively with ddH_2_O, dried in an N_2_-stream, and attached to a flow chamber with double sided tape (2.5 cm x 2 mm x 120 μm) and five min epoxy resin. Each flow chamber was prepared immediately preceding each reaction as follows: 3 min incubation in 1% HBSA (1% BSA in 10 mM Imidazole pH 7.4, 50 mM KCl), 30 sec incubation 4 mg/ml Streptavidin, washed with 1% HBSA, and finally equilibrated with 1x TIRF buffer (20 mM Imidazole pH 7.4, 100 mM KCl, 0.4 mM ATP pH 7.0, 2 mM MgCl_2_, 2 mM EGTA, 20 mM DTT, 30 mM glucose, 0.5% methylcellulose 4000 cP). To initiate reactions, modified actin monomers were rapidly diluted to 1 μM final concentration (containing 0.25-0.5% biotinylated RMA, and 15% Alexa488-RMA, except for Fig. 3 in which Alexa670 labeled R-actin was used) in 1x TIRF buffer supplemented with 2 mg/ml catalase and 10 mg/ml glucose oxidase and transferred into a flow chamber. To measure nucleation, elongation, and branching, the actin-binding proteins of interest were included in the 1xTIRF buffer mix before addition of the actin monomers and transfer into the flow chamber. To measure severing/disassembly, actin monomers were polymerized at room temperature until filaments reached approximately 15 μm in length, then free monomers were washed out and Cof1 was flowed in. Time-lapse TIRF imaging was performed with a Ti2-LAPP inverted microscope equipped with through-the-objective TIRF illumination (Nikon), a LU-4N 4-laser unit (Nikon) and an iXon Ultra 888 EMCCD camera (Andor Technology). The pixel size corresponds to 0.217 μm.

### TIRF analysis

TIRF data was analyzed using ImageJ software. To start, background fluorescence was subtracted using the background subtraction program and a rolling ball radius of 10 pixels. Filament nucleation rates were calculated by counting the number of filaments in a 130 μm x 130 μm box (5 squares per reaction) at 5 minutes after start of the reaction. For each reaction, dilution of actin monomers in the TIRFM buffer was taken as the start of the reaction. Filament elongation rates were determined by measuring the change in filament length over time (at least 60 seconds).

Arp2/3 mediated actin filament branching was measured by counting the total number of branches formed at specific time points (4 to 14 min) after start of the reaction for equally large FOVs (150 x 150 micron). To exclude overlapping filaments, each branch was traced back to its first appearance, and only counted as a branch if it would grow off of another filament for at least 1 min. To account for differences in actin filament nucleation, the branching rate (#/min) for each FOV was determined using an exponential curve fit of at least 5 time points (See Fig. S2). Where needed, more time points were analyzed to obtain a good curve fit (R^2^ = 0.95). The rate constants for each FOV were used for statistical analysis.

Cofilin-mediated filament severing was analyzed by counting the number if severing events observed during the next 600 sec after flow-in of Cof1. To account for differences in the number and length of filaments between FOVs, severing was normalized to the initial length of the filament analyzed, i.e. the length of the filament before flow-in of Cof1. Time to half-maximal severing for each FOV was determined from cumulative severing curves, and used for statistical analysis. Best curve fit (R^2^ ≥ 0.97) was obtained using the equation for specific binding with Hill slope.

### Statistical analysis

Statistical analysis, and curve fitting was performed using Prism 5.0. The specific curve fitting algorithms used for each experiment are described above in TIRF analysis. The distribution of the rate constants obtained from the curve fits for each FOV are represented as bee swarms, overlaid with box-and whisker plots that show minimum, median, and maximum. Statistical difference between the distributions of two differently modified actins was analyzed using a two-tailed unpaired t-test.

## Legends Supplemental Figures

**Figure S1**

**(A)** Still images showing the difference in nucleation between the Ac-, R-, M-, and D-actin at 5 minutes after initiating the reaction. **(B)** Montages of representative TIRF microscopy movies demonstrating elongation of different actins in the presence of 10 nM mDia1 and 1.5 μM PFN. Arrowheads indicate filament ends.

**Figure S2**

**(A)** Distribution of the number (#) of branches that are formed 8 min after start of the reaction (N ≥ 15, 3 replicates with 5 FOV each). Corresponding box and whisker plots are shown as overlays. Statistical significance was tested by comparing two different actins using an unpaired t-test. **(B)** Graphs showing the individual branching curves (including fit) for each FOV of the different reactions performed with the differently modified actins. These curves were used to determine the branching rates represented in Fig 2C.

**Figure S2**

Graphs show the individual cumulative severing curves (including fit) for each FOV of the different reactions performed with the differently modified actins. The average and StDev of these curves for each actin is shown in Fig 4B. The time to half-maximal severing of each individual curve is represented in Fig 4C.

