## Supplementals for "N-terminal modification of actin by acetylation and arginylation determines the architecture and assembly rate of linear and branched actin networks"

### Slide 1
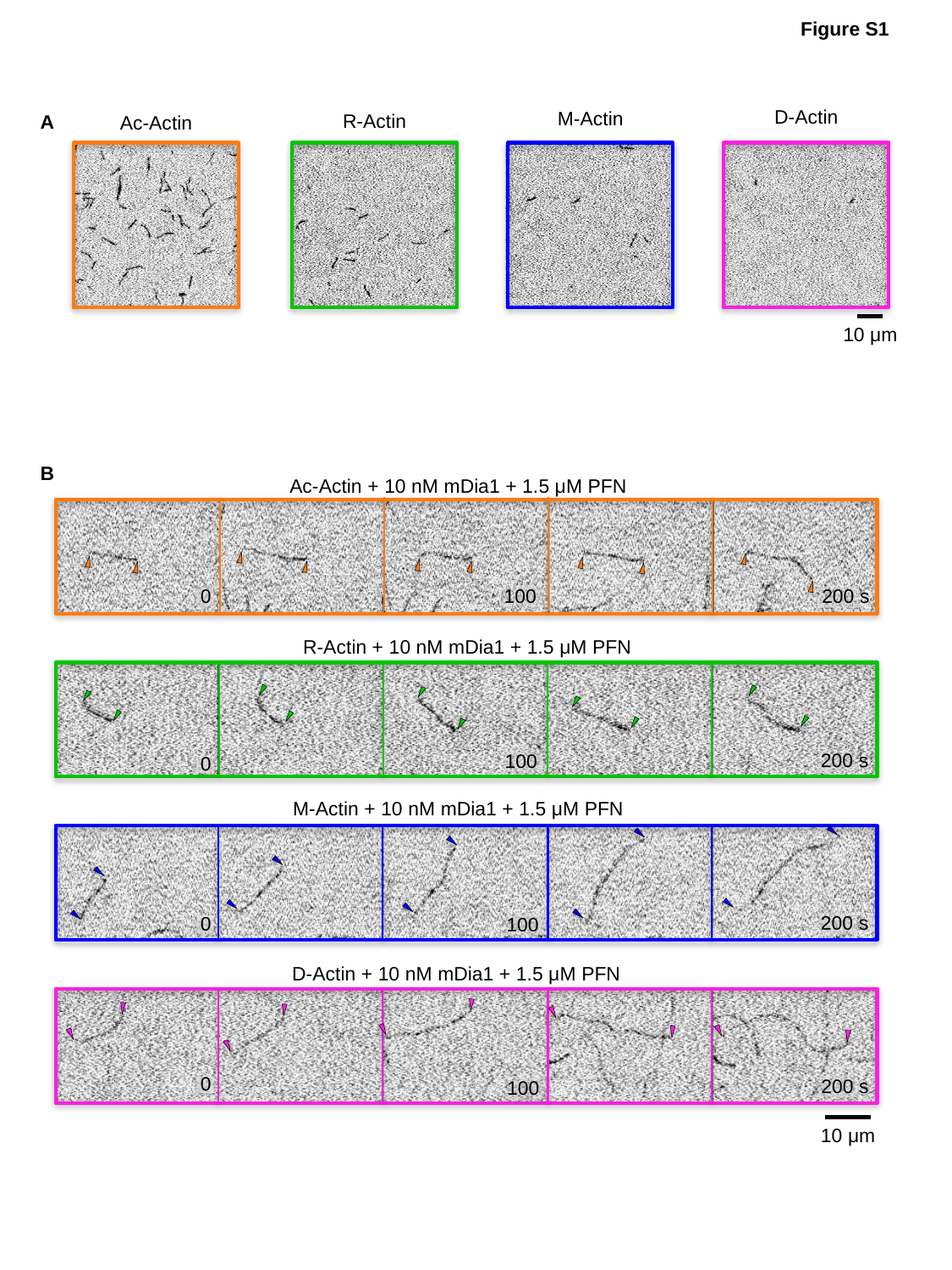

Figure S1
D-Actin
M-Actin
R-Actin
A
Ac-Actin
10 μm
B
Ac-Actin + 10 nM mDia1 + 1.5 μM PFN
0
100
200 s
R-Actin + 10 nM mDia1 + 1.5 μM PFN
200 s
100
0
M-Actin + 10 nM mDia1 + 1.5 μM PFN
200 s
0
100
D-Actin + 10 nM mDia1 + 1.5 μM PFN
0
200 s
100
10 μm

### Slide 2
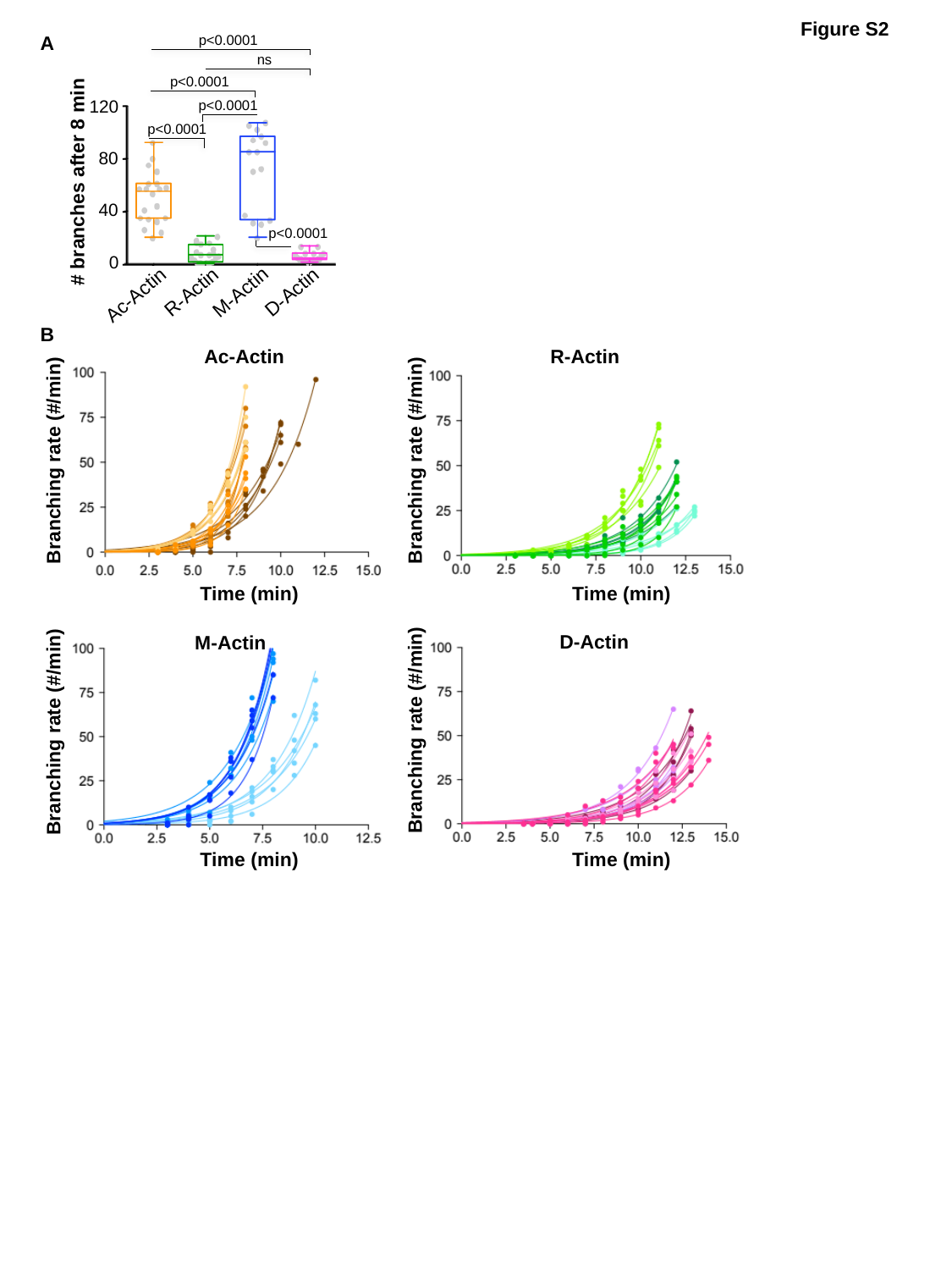

Figure S2
p<0.0001
A
ns
p<0.0001
120
p<0.0001
p<0.0001
80
#### branches after 8 min
40
p<0.0001
0
R-Actin
D-Actin
M-Actin
Ac-Actin
B
Ac-Actin
R-Actin
Branching rate (#/min)
Branching rate (#/min)
Time (min)
Time (min)
D-Actin
M-Actin
Branching rate (#/min)
Branching rate (#/min)
Time (min)
Time (min)

### Slide 3
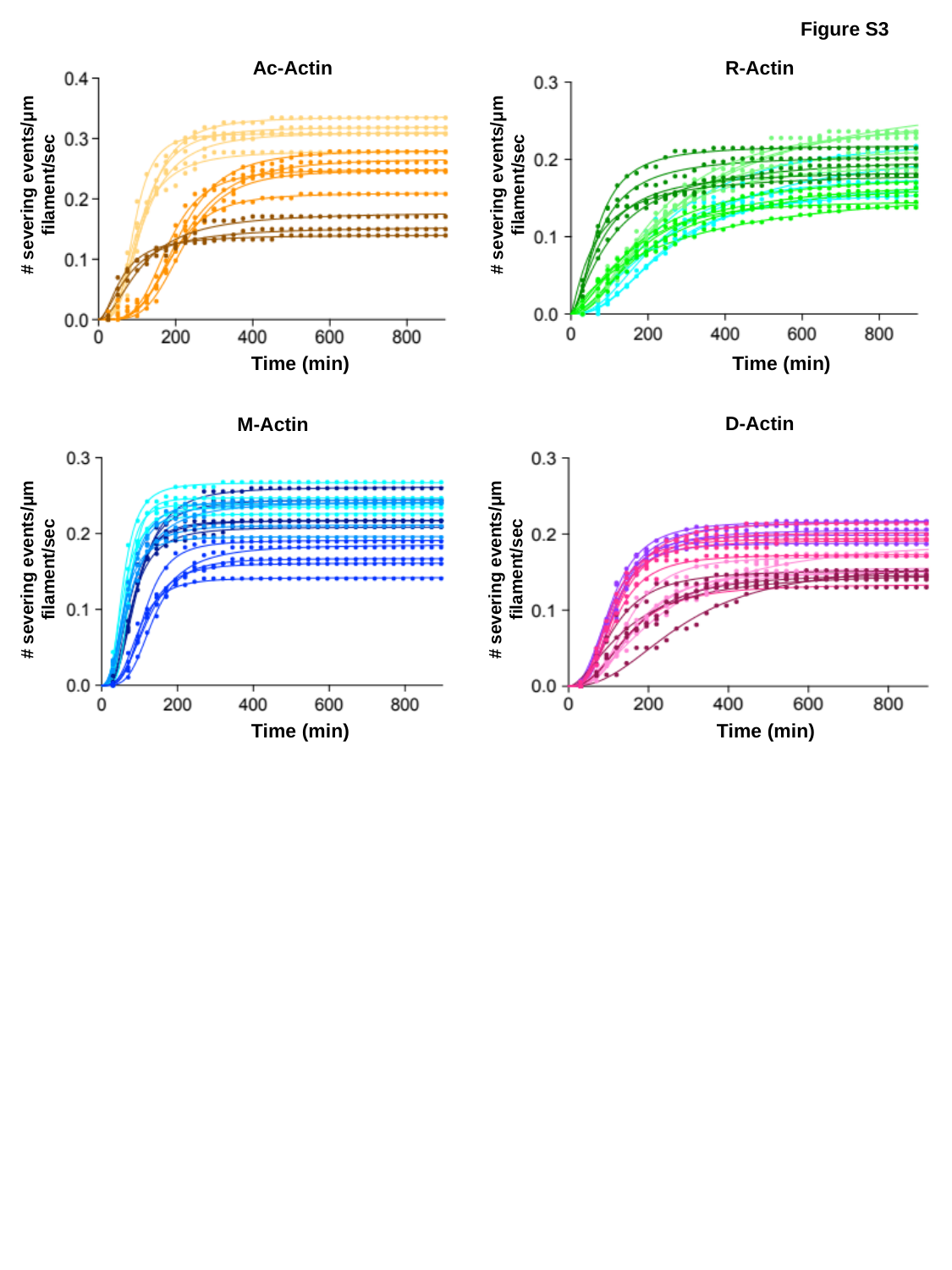

Figure S3
Ac-Actin
R-Actin
#### severing events/μm filament/sec
#### severing events/μm filament/sec
Time (min)
Time (min)
D-Actin
M-Actin
#### severing events/μm filament/sec
#### severing events/μm filament/sec
Time (min)
Time (min)
